# The landscape of micro-inversions provides clues for population genetic analysis of humans

**DOI:** 10.1101/2020.07.27.218867

**Authors:** Li Qu, Luotong Wang, Feifei He, Yilun Han, Longshu Yang, May D. Wang, Huaiqiu Zhu

## Abstract

**Background:** Variations in the human genome have been studied extensively. However, little is known about the role of micro-inversions (MIs), generally defined as small (<100 bp) inversions, in human evolution, diversity, and health. Depicting the pattern of MIs among diverse populations is critical for interpreting human evolutionary history and obtaining insight into genetic diseases.

**Results:** In this paper, we explored the distribution of MIs in genomes from 26 human populations and 7 nonhuman primate genomes and analyzed the phylogenetic structure of the 26 human populations based on the MIs. We further investigated the functions of the MIs located within genes associated with human health. With hg19 as the reference genome, we detected 6,968 MIs among the 1,937 human samples and 24,476 MIs among the 7 nonhuman primate genomes. The analyses of MIs in human genomes showed that the MIs were rarely located in exonic regions. Nonhuman primates and human populations shared only 82 inverted alleles, and Africans had the most inverted alleles in common with nonhuman primates, which was consistent with the “Out of Africa” hypothesis. The clustering of MIs among the human populations also coincided with human migration history and ancestral lineages.

**Conclusions:** We propose that MIs are potential evolutionary markers for investigating population dynamics. Our results revealed the diversity of MIs in human populations and showed that they are essential to constructing human population relationships and have a potential effect on human health.

## 1 Background

As a kind of structural variation (SV) in genomes, inversions are defined as chromosome rearrangements in which a segment of a chromosome is reversed end to end [1]. Usually, an inversion occurs when a single chromosome undergoes breakage and rearrangement within itself. Although it has long been known that inversions are associated with primate evolution [2], they were only recently found to play an important role in human evolution and diseases [3,4]. A number of studies have focused on inversions in the human genome [5, 6], and for decades, many detectable macroinversion polymorphisms in humans have been verified by experiments and proved to have been significant in human evolutionary history [7,8]. For example, inversions located in 8p23.1 have been reported to be associated with autoimmune and cardiovascular diseases [3]. In addition, inversions have been designated as evolutionary markers of human phenotypic diversity [9,10]. For example, a common 900 kb inversion polymorphism at 17q21.31 associated with Parkinson’s disease suggested multiple distributions of inversions among ethnic populations [7]. Indeed, with the development of inversion detection methods, inversions are being increasingly recognized as one of the most important mechanisms underlying genetic diversity.

However, the studies mentioned above were mainly limited to the detection of large-scale inversions, usually >100 kb in length. Recently, small-scale inversions (with lengths much shorter than 10 kb) have been used in several studies for purposes such as phylogenetic inference. These studies found an influence of small inversions on the formation of unusual flanking sequences in human and chimpanzee genomes [11] and developed tools to detect in-place inversions [12]. Nevertheless, the results of these studies were highly variable because of differences in the size used to define small inversions [13]. Moreover, current studies focus mainly on the differences in small inversions between human (*Homo sapiens*) and chimpanzee (*Pan troglodytes*) genomes [11], excluding the genomes of more distantly related primates such as gorillas (*Gorilla*), orangutans (*Pongo pygmaeus*), gibbons (*Hylobates sp.*), baboons (*Papio anubis*), and rhesus macaques (*Macaca mulatta*).

Among all the inversions of small size, micro-inversions (MIs) [13], which are a type of extremely small inversion (generally <100 bp) found remarkably in the human genome, have an uncertain function in the research community. However, statistical analysis has suggested that MIs, which are usually similar to other rare genomic changes, may serve as phylogenetic markers because of their rare occurrence and low homoplasy [14]. Furthermore, many studies have examined the functions of inversions among multiple nonprimate species, such as yeast, sticklebacks, grasshoppers, *Drosophila*, ducks, chicken, and mice [15-21]. Regarding primates, which have larger genomes, comparative studies of the influence of chromosomal inversions occurring within chromosomes in the human and chimpanzee genomes can be traced back to the last century [2]. Nevertheless, such studies on human inversions are rare because of the limitations of detection techniques [22]. Great efforts have been focused on developing experimental techniques to discover large-scale inversions, including methods based on single-cell sequencing [23, 24], read assembly [25], and probe hybridization and amplification [26]. However, these experimental methods are usually expensive, time-consuming, and not aimed at detecting small inversions. With the advent of next-generation sequencing (NGS), the 1000 Genomes Project (1KGP) has provided a large amount of whole-genome sequencing data for individuals from a variety of populations across the world [27,28]. However, the short reads generated by NGS in the 1KGP are generally very short (<100 bp), making both the detection and analysis of MIs difficult. In contrast to well-studied large inversions, which cause more abnormal short reads and thus are discovered more frequently based on NGS short reads, MIs have not yet been studied because it is difficult to detect MIs shorter than the read length, with most identified as unmapped reads and thus discarded before further analysis. However, a large proportion of the sequenced short reads in the 1KGP are unmapped short reads that may contain SVs, many of which could be MIs. For example, in NA18525, a low-coverage sequencing sample in the 1KGP, 608,982,899 short reads are mapped reads, while 360,144,413 (37.2% of the total) are unmapped reads. The Micro-inversion Detection (MID) method developed in our previous study [13] uses these unmapped reads to detect MIs with good performance. The algorithm of MID was designed based on a dynamic programming path-finding approach, which can efficiently and reliably identify MIs from unmapped short NGS reads. This process subsequently facilitates the analysis of MIs across many populations based on high-throughput sequencing data. Therefore, an increased understanding of the MI landscape across various human populations will facilitate comparative analyses among individuals by taking individual differences into account.

Although efforts have been made to analyze small inversions that are still >100 bp in nonhuman organisms [11,12], comprehensive studies of the effects of MIs, which are < 100 bp in length, on human diversity, evolution, and diseases in a large number of human genomes in the 1KGP are lacking. In this study, we set out to detect MIs and further investigate the roles of MIs in the diversity and evolution of 26 human populations and 7 nonhuman primate populations. Overall, we explored the distribution of MIs in all 26 populations from the 1KGP [27] and detected 6,968 MIs within all 1,937 human samples and 24,476 MIs in 7 nonhuman primate genomes. Furthermore, we analyzed the extent of MI diversity and built phylogenetic trees by inverted allele counts at both the population and species scales. These results indicated that MIs rarely occur in or near protein-coding genes and that only a few are common in both primate and human populations. They also show that Africans share the most inverted alleles with nonhuman primates, which is consistent with the “Out of Africa” hypothesis [29]. More importantly, the phylogenetic analysis demonstrates that MIs are sensitive evolutionary markers for categorizing all populations. The categories coincide with human migration history and ancestral lineages. The analysis also revealed possible functions of some MIs located within disease-causing genes. Thus, MIs should be considered for studies of human evolution.

## 2 Results

### 2.1 Overview and distribution of MIs

Our previous work [13] focused on the detection of MIs in the initial phases of the 1KGP, which encompassed fewer individuals from only 19 populations and was largely limited to the statistical analysis of the MI distribution and polymorphisms, despite the few disease-related genes available for analysis. Herein, we constructed a more integrated map of MIs, including extra and new samples from the 1KGP phase III data. In addition, compared with our previous work, the current study presents a more comprehensive analysis that emphasizes the population diversity of MIs, quantifies the genetic impact of MIs, and explores the important roles that MIs play in human evolution. The flowchart of the entire analysis presented in this study is shown in Fig. 1.

**Fig. 1.**
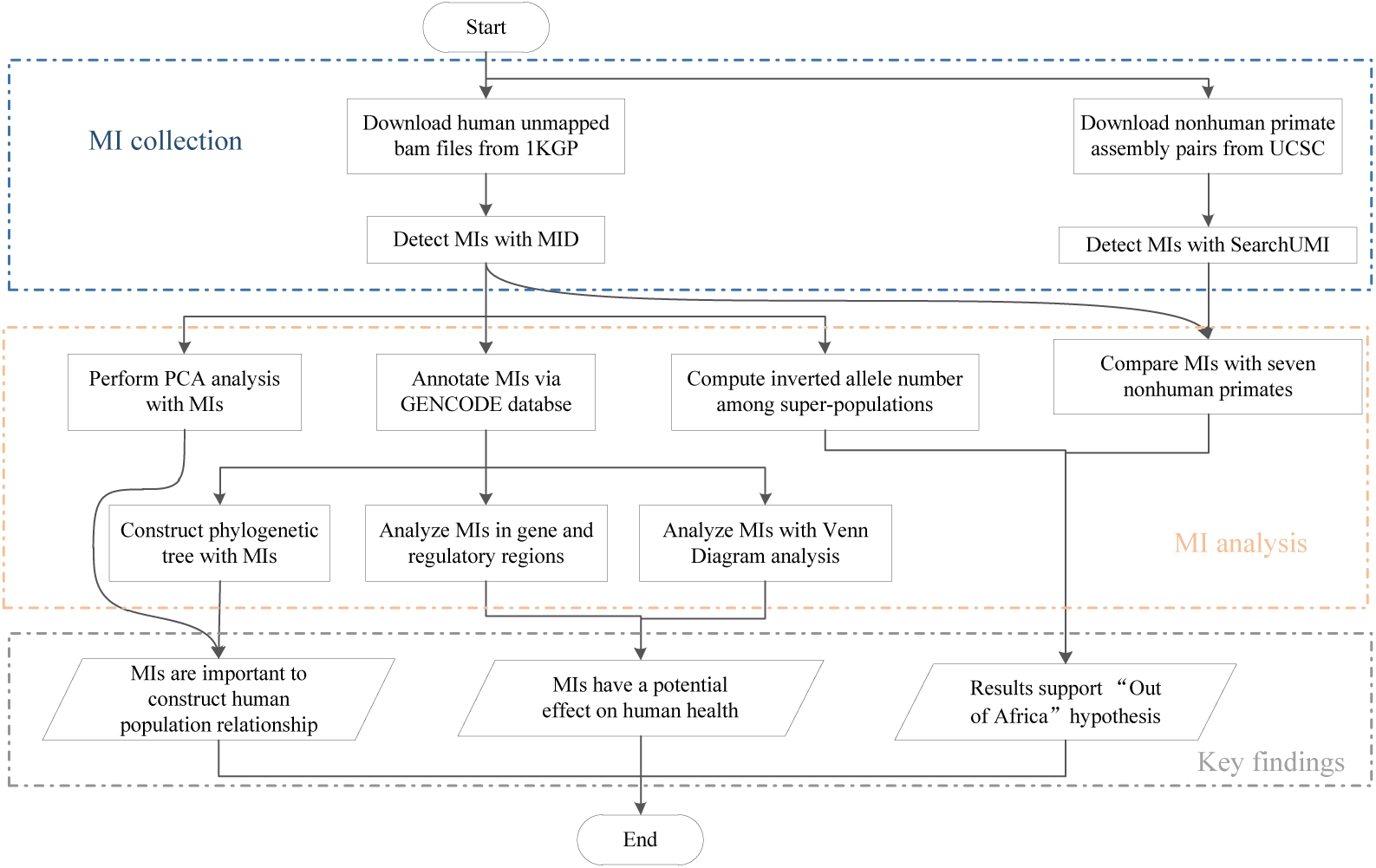
A flowchart outlining the procedures of MI calling, MI analysis, and the key findings.

Although the recurrence of specific MIs is rare, a series of MIs often converge to particular regions in the genome. To provide a more intuitive analysis of MIs, by extending the boundaries of overlapping MIs, we defined MI regions (MIRs) similar to the copy number variation (CNV) regions proposed by Yang *et al* [30]. Thus, MIRs refer to the united regions of overlapping MIs, with the requirement that an MIR must be at most four base pairs longer than any one of its included MIs (shown in Fig. 2A). According to the definition of an MIR, each MI can be contained in only one MIR, and each MIR may contain at least one MI. Thus, the MIs in the same MIR have almost the same start and end positions on the chromosome. It should be noted that MIs merged within the same MIR in our article could not originate from the same individual because MIs with the same end limits detected in the same individual were merged into one MI during the MID detection process [13]. Furthermore, the MIs merging in the same MIR were those in different individuals at the same region.

**Fig. 2.**
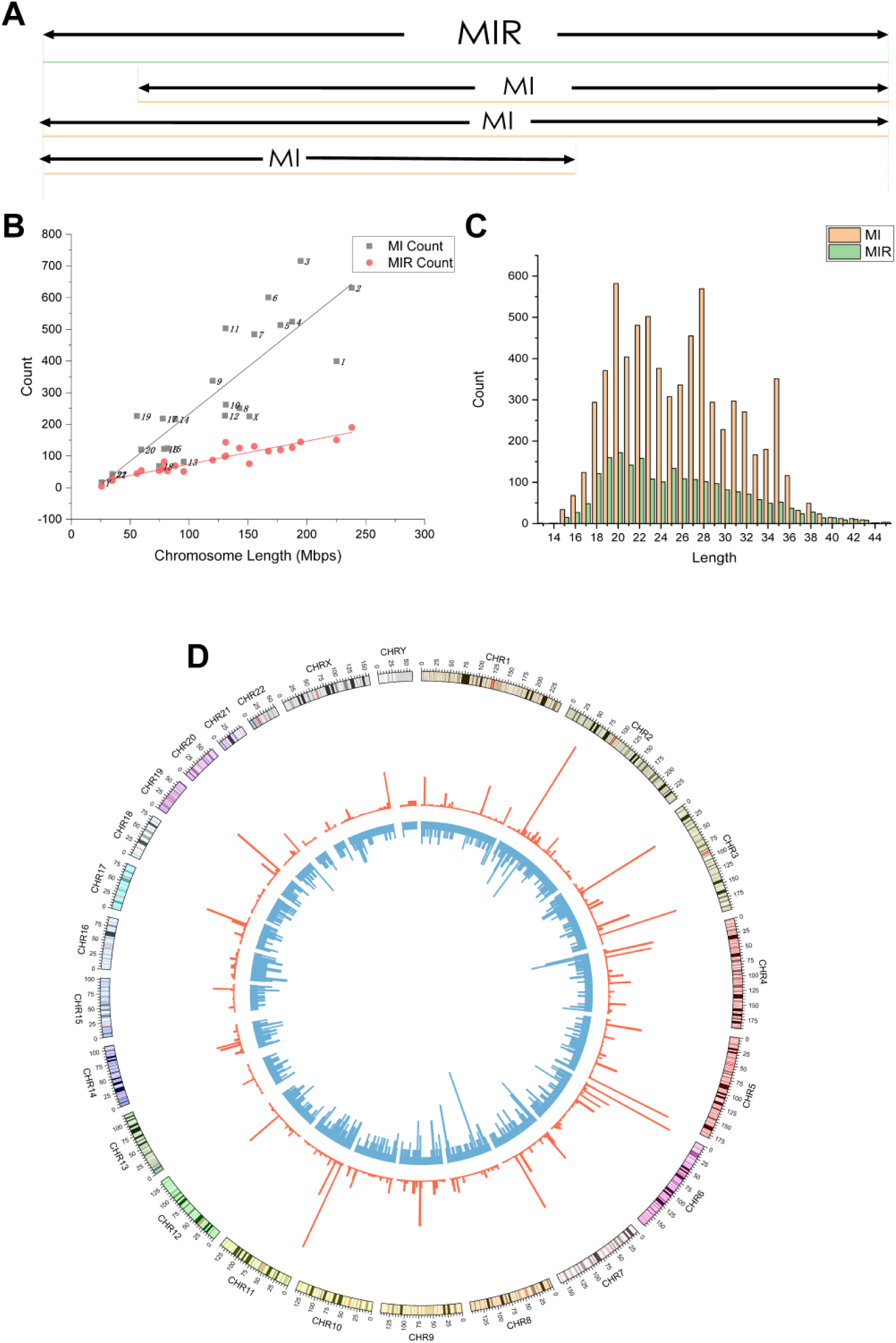
MIs and MIRs in human chromosomes. (A) Schematic showing how an MIR is a region where MIs overlap (black bars, MIRs; blue bars, MIs in one individual). (B) Scatter plot of MIR count against chromosome length. (C) Length distribution of 6,968 MIs (13-45 bp) and 2,140 MIRs (14-45 bp). (D) Distribution of locations of MIs on chromosomes.

Since the MI could be considered a polymorphic variant that exists at a given position in the genome, one MI has two alleles. In this study, we refer to the allele found in the reference genome as the reference allele and the allele with the opposite inverted sequence against the reference genome as the inverted allele. We also refer to the inverted allele found in the common ancestor of the 7 nonhuman primates and the humans as the ancestral allele and to the inverted allele, which appeared by mutation at some point over time, as the derived allele.

In this study, using the MID method [13], we identified 6,968 MIs (see Table 1 for a list and statistics) among all unmapped reads in 1,937 samples from the 26 human populations, which were merged into 2,140 MIRs (the full list of sample names is provided in Table S1). Among the 6,968 MIs, five occurred in more than 200 individual genomes. In addition, among the 2,140 MIRs, 1,169 (54.6%) overlapped with gene regions, among which 1,063 (90.9%) overlapped with intronic and the remaining 106 overlapped with exonic regions (see Tables S2-S4 for the full list of MIs, MIRs, and MIs in exonic regions, respectively). Of the 106 MIRs in exonic regions, 30 (28.3%) overlapped with coding sequence (CDS) regions, 39 (36.8%) with untranslated regions (UTRs), and 37 (34.9%) with other functional regions, including miRNA and mt_rRNA regions. Of the 971 MIRs in intergenic regions, 47 MIRs (4.8%) overlapped with candidate promoter regions.

**Table 1.**
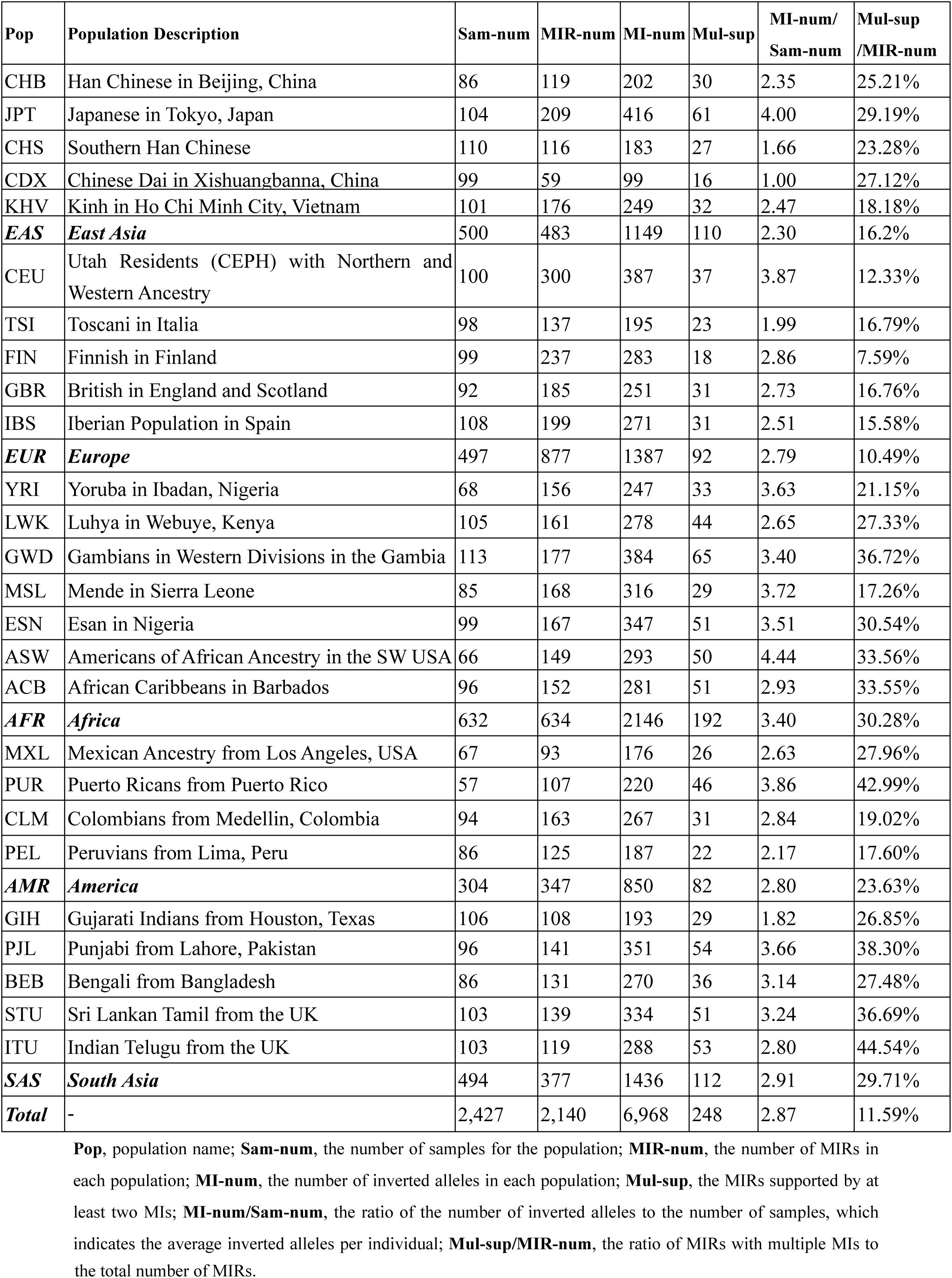
Overview of MIs detected in 1,937 samples.

| Pop | Population Description | Sam-num | MIR-num | MI-num | Mul-sup | MI-num/<br>Sam-num | Mul-sup<br>/MIR-num |
| --- | --- | --- | --- | --- | --- | --- | --- |
| CHB | Han Chinese in Beijing, China | 86 | 119 | 202 | 30 | 2.35 | 25.21% |
| JPT | Japanese in Tokyo, Japan | 104 | 209 | 416 | 61 | 4.00 | 29.19% |
| CHS | Southern Han Chinese | 110 | 116 | 183 | 27 | 1.66 | 23.28% |
| CDX | Chinese Dai in Xishuangbanna, China | 99 | 59 | 99 | 16 | 1.00 | 27.12% |
| KHV | Kinh in Ho Chi Minh City, Vietnam | 101 | 176 | 249 | 32 | 2.47 | 18.18% |
| **EAS** | **East Asia** | 500 | 483 | 1149 | 110 | 2.30 | 16.2% |
| CEU | Utah Residents (CEPH) with Northern and Western Ancestry | 100 | 300 | 387 | 37 | 3.87 | 12.33% |
| TSI | Toscani in Italia | 98 | 137 | 195 | 23 | 1.99 | 16.79% |
| FIN | Finnish in Finland | 99 | 237 | 283 | 18 | 2.86 | 7.59% |
| GBR | British in England and Scotland | 92 | 185 | 251 | 31 | 2.73 | 16.76% |
| IBS | Iberian Population in Spain | 108 | 199 | 271 | 31 | 2.51 | 15.58% |
| **EUR** | **Europe** | 497 | 877 | 1387 | 92 | 2.79 | 10.49% |
| YRI | Yoruba in Ibadan, Nigeria | 68 | 156 | 247 | 33 | 3.63 | 21.15% |
| LWK | Luhya in Webuye, Kenya | 105 | 161 | 278 | 44 | 2.65 | 27.33% |
| GWD | Gambians in Western Divisions in the Gambia | 113 | 177 | 384 | 65 | 3.40 | 36.72% |
| MSL | Mende in Sierra Leone | 85 | 168 | 316 | 29 | 3.72 | 17.26% |
| ESN | Esan in Nigeria | 99 | 167 | 347 | 51 | 3.51 | 30.54% |
| ASW | Americans of African Ancestry in the SW USA | 66 | 149 | 293 | 50 | 4.44 | 33.56% |
| ACB | African Caribbeans in Barbados | 96 | 152 | 281 | 51 | 2.93 | 33.55% |
| **AFR** | **Africa** | 632 | 634 | 2146 | 192 | 3.40 | 30.28% |
| MXL | Mexican Ancestry from Los Angeles, USA | 67 | 93 | 176 | 26 | 2.63 | 27.96% |
| PUR | Puerto Ricans from Puerto Rico | 57 | 107 | 220 | 46 | 3.86 | 42.99% |
| CLM | Colombians from Medellin, Colombia | 94 | 163 | 267 | 31 | 2.84 | 19.02% |
| PEL | Peruvians from Lima, Peru | 86 | 125 | 187 | 22 | 2.17 | 17.60% |
| **AMR** | **America** | 304 | 347 | 850 | 82 | 2.80 | 23.63% |
| GIH | Gujarati Indians from Houston, Texas | 106 | 108 | 193 | 29 | 1.82 | 26.85% |
| PJL | Punjabi from Lahore, Pakistan | 96 | 141 | 351 | 54 | 3.66 | 38.30% |
| BEB | Bengali from Bangladesh | 86 | 131 | 270 | 36 | 3.14 | 27.48% |
| STU | Sri Lankan Tamil from the UK | 103 | 139 | 334 | 51 | 3.24 | 36.69% |
| ITU | Indian Telugu from the UK | 103 | 119 | 288 | 53 | 2.80 | 44.54% |
| **SAS** | **South Asia** | 494 | 377 | 1436 | 112 | 2.91 | 29.71% |
| **Total** | - | 2,427 | 2,140 | 6,968 | 248 | 2.87 | 11.59% |

**Pop**, population name; **Sam-num**, the number of samples for the population; **MIR-num**, the number of MIRs in each population; **MI-num**, the number of inverted alleles in each population; **Mul-sup**, the MIRs supported by at least two MIs; **MI-num/Sam-num**, the ratio of the number of inverted alleles to the number of samples, which indicates the average inverted alleles per individual; **Mul-sup/MIR-num**, the ratio of MIRs with multiple MIs to
the total number of MIRs.

Information regarding the 30 MIs overlapping with coding sequence (CDS) regions is provided in Table S5. As shown in Table S5, most of the 30 MIs were located within CNV loss or gain regions. This result indicates that the MIs may affect gene function combined with other kinds of SVs. Additionally, 10 of the 30 MIs include Single nucleotide polymorphisms (SNPs) according to the annotation of the dbSNP [31] or the Exome Aggregation Consortium (ExAC) [32] database. The bases marked in red color in the column “Reference allele sequence” and “Inverted allele sequence” are the SNP polymorphism positions contained in the MI sequences. It should be noted that the inverted allele of chr11: 126146988-126147016 included the SNP at position 126147003 (A>G) named rs7116126 in the dbSNP database, which was annotated by the ClinVar [33] database as indicating a conflicting interpretation of pathogenicity. Thus, the pathogenicity of this SNP allele is not clear, and various studies have reported conflicting interpretations of this SNP. To further analyze this MI polymorphism, we listed the details of this MI annotated by multiple databases from the UCSC genome browser (http://genome.ucsc.edu/) [34] in Fig. S4, showing that the MI region is conserved in a few vertebrates such as the human, rhesus macaque, mouse, dog, elephant, and chicken, which indicates that this region may not change easily. Further evidence is needed to determine the specific function of this MI linked with a few SVs and SNP rs7116126.

A scatter plot of the inverted allele number and MIR count against chromosome length is shown in Fig. 2B. Generally, the numbers of inverted alleles and MIRs are positively correlated with the lengths of chromosomes (Fig. 2B), which is reasonable, because longer chromosomes have more DNA bases and thus a greater chance of DNA replication errors. Additionally, the correlations between MI and MIR counts and length were analyzed using Pearson’s correlation analysis. Specifically, a strong correlation was detected between the MIR count and chromosome length, where the Pearson’s correlation coefficient was r = 0.935 (P<0.0001); the corresponding result for MI was 0.861 (P<0.0001). A scatter plot of MI and MIR counts against gene density is shown in Fig. S1. Although we found no strong linear correlation between the number of inverted alleles and the gene density, we found a phenomenon in which most scatter above the line of the best fit in Fig. 2B, which was attributable to chromosomes with high gene densities. For example, we discovered that chromosomes 19, 17, 22, and 11, which were responsible for the scatter above the line of best fit, have high gene density, as shown in Fig. S1. This suggests that gene density may also affect the inverted alleles of a chromosome, while this relationship is not strictly linear. The MI and MIR event rate distribution across chromosomes is shown in Fig. S2. As shown in Fig. 2C, the lengths of the MIs varied from 15 to 45 bp but were concentrated within 18 to 30 bp. The specific positions of MIs among 24 chromosomes are shown in Fig. 2D. Notably, when comparing the MIs in our study with the inversion-calling results from the phase III analysis of the 1KGP, we found that the shortest inversion detected by the 1KGP was 257 bp, which was much longer than 100 bp. We also noted that in other studies on inversions, no MIs in the range from 10 to 100 bp were discussed. This indicates a discrepancy in length between the MIs in this study and traditional inversions in previous studies. Thus, our analysis of MIs (<100 bp), overlooked by all the studies including 1KGP analysis, sheds light on the limited understanding of human SVs.

### 2.2 Inverted allele number per individual among populations

SNPs and SVs in the 1KGP have been reported to display variant polymorphism and reveal genetic diversity among 26 populations [27]. However, MIs, as a kind of SVs, have never been studied to capture the genetic diversity among 26 populations. Thus, we report an analysis of MI diversity in this subsection.

We calculated an average of 2.87 inverted alleles per individual for all individuals. A bar plot showing the average counts of inverted alleles per individual for five super-populations (African, American, South Asian, East Asian, and European) is presented in Fig. S3, which shows that the African super-population had the most inverted alleles per individual at 3.4, while the East Asian population had the fewest at 2.3. As we focused more on the effect of inverted alleles instead of reference alleles on evolution, we also display the average count of inverted alleles in the five super-populations, without including the 490 samples with no MIs detected, in Fig. 3A. This analysis showed that when considering only individuals with the inverted alleles, the five super-populations ranked in descending order according to the average count of inverted alleles per individual as follows: African > American > European > South Asian > East Asian. Notably, the African super-population had the most inverted alleles per individual at 3.96, while the East Asian population had the fewest at 3.03. The results displayed in both Fig. S3 and Fig. 3A are consistent with those of recent studies employing human SNP analysis [27,35], in which the African super-population has the largest SNP number and the East Asian super-population has the smallest SNP number. In addition, a phylogenetic tree was constructed according to the MIs in gene regions for the five super-populations (see Fig. 3B). The result implies that the African super-population is distinct from the other 4 super-populations. The descending order of the super-populations based on average inverted allele number and the clustering result also fit the Out of Africa hypothesis [29], which posits that humans originated in Africa and then modern Africans migrated to other continents. We will discuss this point further in the Discussion and conclusions section. Regarding the 26 populations, the average inverted allele count per individual populations ranged from 1.00 to 4.44 (see Table 1). Furthermore, the population of Americans of African Ancestry in the SW USA (ASW) had the most inverted alleles at 4.44, and the Chinese Dai in Xishuangbanna, China (CDX), had the least at 1.00. The geographic locations of the 26 populations are shown in Fig. 3C.

**Fig. 3.**
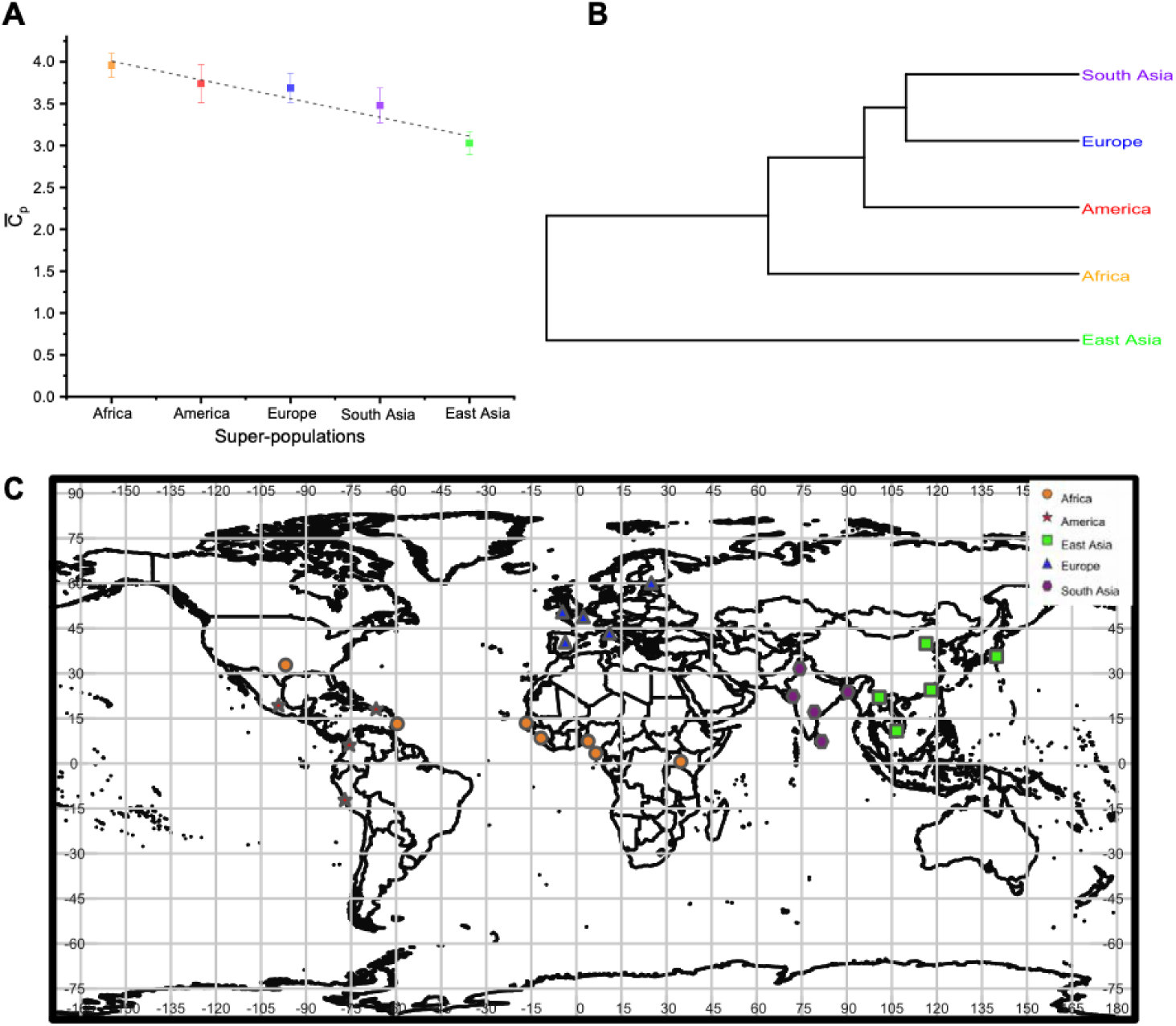
Average counts of inverted alleles per individual. 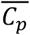 is the average count of inverted alleles per individual. Different colors represent the five super-populations: orange, African; red, American; blue, European; purple, South Asian; and green, East Asian. (A) Bar plot of the average count of inverted alleles per individual among the five super-populations. The data are presented as the mean ± standard error (SE). The average count ranged from 3.03 to 3.96. (B) Phylogenetic tree based on the average pairwise differences between super-populations according to the MIs in gene regions. (C) Geographic locations of the 26 populations.

### 2.3 Population structure based on MI statistics

To examine the genetic structure of the MIs hierarchically among populations, we constructed a genetic distance-based phylogenetic tree using all the MIs detected from the 26 populations (Fig. 4A), which yielded results that generally provided evidence of genetic clustering: African and American populations were clustered on one branch, European and South Asian populations were clustered on a second branch, and East Asian populations were clustered on a third branch. Since the gene regions are generally considered as conserved domains, the MIs therein may reflect the human evolution and population relationship better than in intergenic regions. Therefore, we also constructed a phylogenetic tree with only MIs in gene regions (Fig. 4B). Similar to the analysis of all MIs presented in Fig. 4A, the phylogenetic analysis of MIs in gene regions presented in Fig. 4B clustered the populations into four branches. The topology of the tree not only met our expectation but also reflected a regional pattern among the 26 populations, suggesting that population groups that live geographically closer to one another have shorter MI-based genetic distances. An exception was the Finnish in Finland (FIN) population (clustered with the East Asian populations in Fig. 4A and forming a single cluster in Fig. 4B), which deviated from the European branch. This deviation of FIN is understandable because Finland is genetically distant from the rest of the European populations [36]. In addition, the Finnish are treated as a separate population rather than a European population in some studies [36]. The phylogenetic tree revealed that MIs with functions similar to those of SNPs are key to tracing the evolution of human population genetic structure. The matrices for constructing the phylogenetic trees presented in Fig. 4A and Fig. 4B are shown in Tables S6 and S7, respectively.

**Fig. 4.**
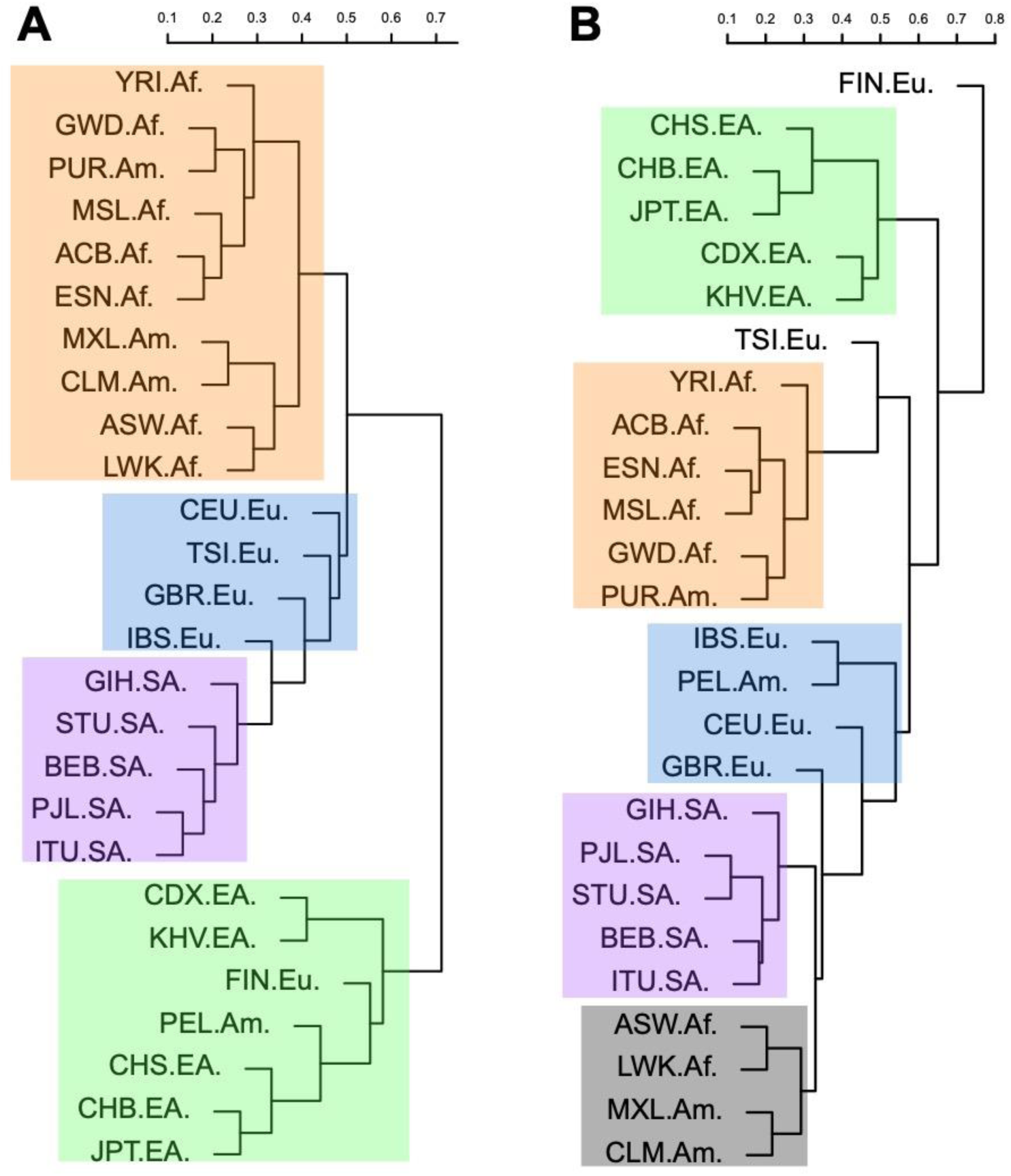
Phylogenetic trees of 26 populations constructed by neighbor joining. (A) Tree for the 26 populations based on average pairwise population distances calculated with all 6,968 MIs. (B) Tree based on average pairwise population distances calculated with the 2,135 MIs in gene regions.

To further reveal undiscovered MI-based relationships among the 26 populations, we performed principal component analysis (PCA) of the 26 populations with all 6,968 MIs. As shown in Fig. 5, the PCA of the MI patterns divided the 26 populations into four groups according to the top two main components. This result indicated that populations in the same super-population were closer to each other than populations in different super-populations. We found that African, European, South Asian, and East Asian populations were clearly recognizable, which represented evolution via genetic drift or other factors. In contrast to those in the other four super-populations, the populations in the American super-population were dispersed and did not form a distinct cluster. Our results are highly consistent with the widespread pattern observed with PCA based on SVs [27].

**Fig. 5.**
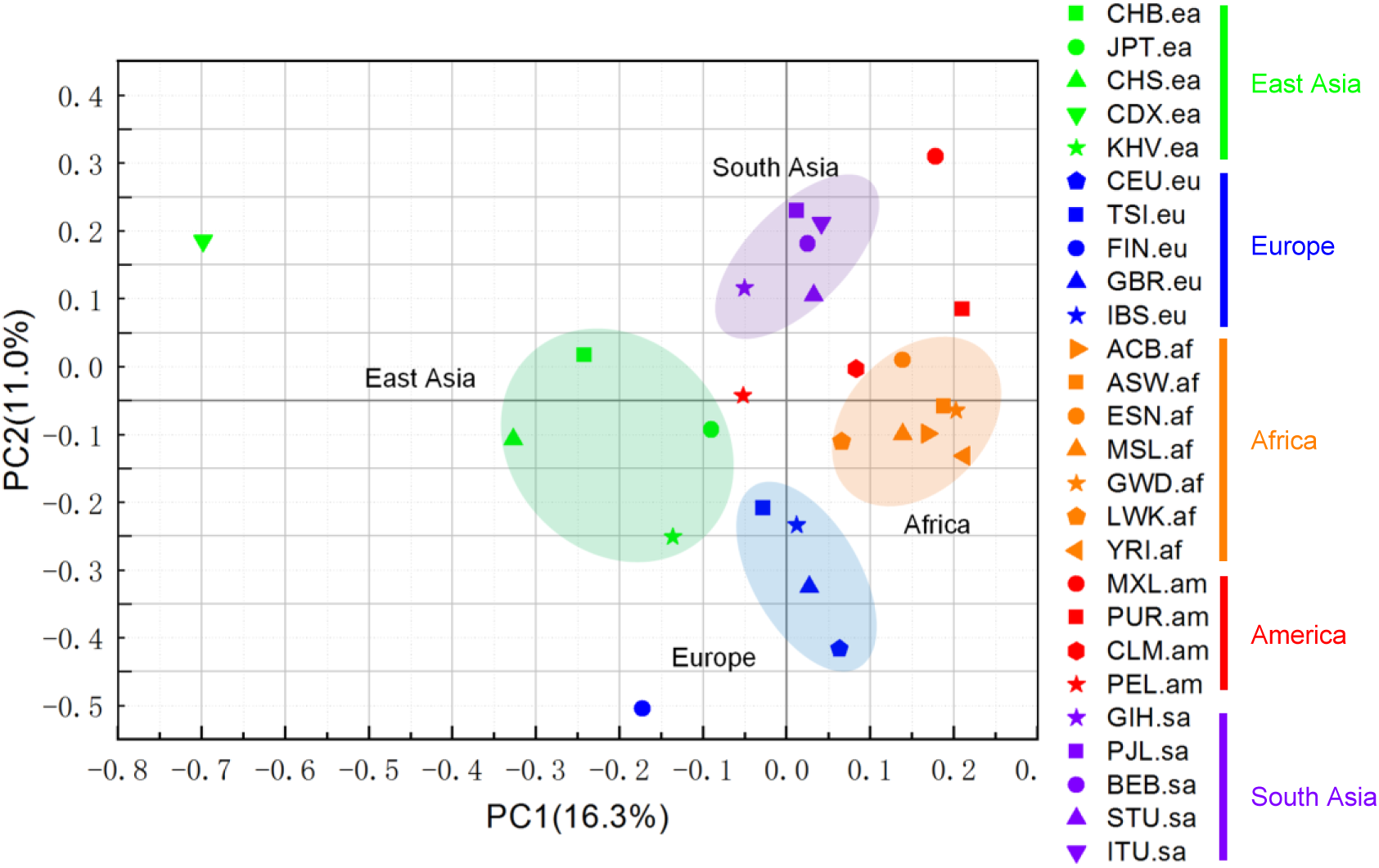
PCA of 26 populations. The first and second principal components are plotted. Different color represent the five super-populations: orange, African; red, American; blue, European; purple, South Asian; and green, East Asian.

### 2.4 Analysis of MIR sharing among the five super-populations

Since the distinct variation pattern of each population may implicitly affect the phenotypic divergence among the populations, it is important to explore the common and distinct MIRs among the five super-populations to determine whether there are differences in MIs among the super-populations. Therefore, to further investigate the diversity and relationships of MIs among the five super-populations, we conducted MIR sharing analysis with a Venn diagram by counting all the MIRs (Fig. 6A). We found that 76 MIRs were shared by all five super-populations. Furthermore, we used only the MIRs located in gene regions to construct the Venn diagram presented in Fig. 6B. A total of 34 MIRs within gene regions annotated by the GENCODE database was shared by all five super-populations (Fig. 6B). In addition, no MIRs in gene regions were shared by the American, European, South Asian, and East Asian super-populations, except for the 34 MIRs that were shared by all five super-populations. However, any four of the super-populations, as long as Africa was included, shared at least one MIR in addition to the 34. This result suggests that the American, European, South Asian, and East Asian super-populations are more closely related to the African super-population than to each other.

**Fig. 6.**
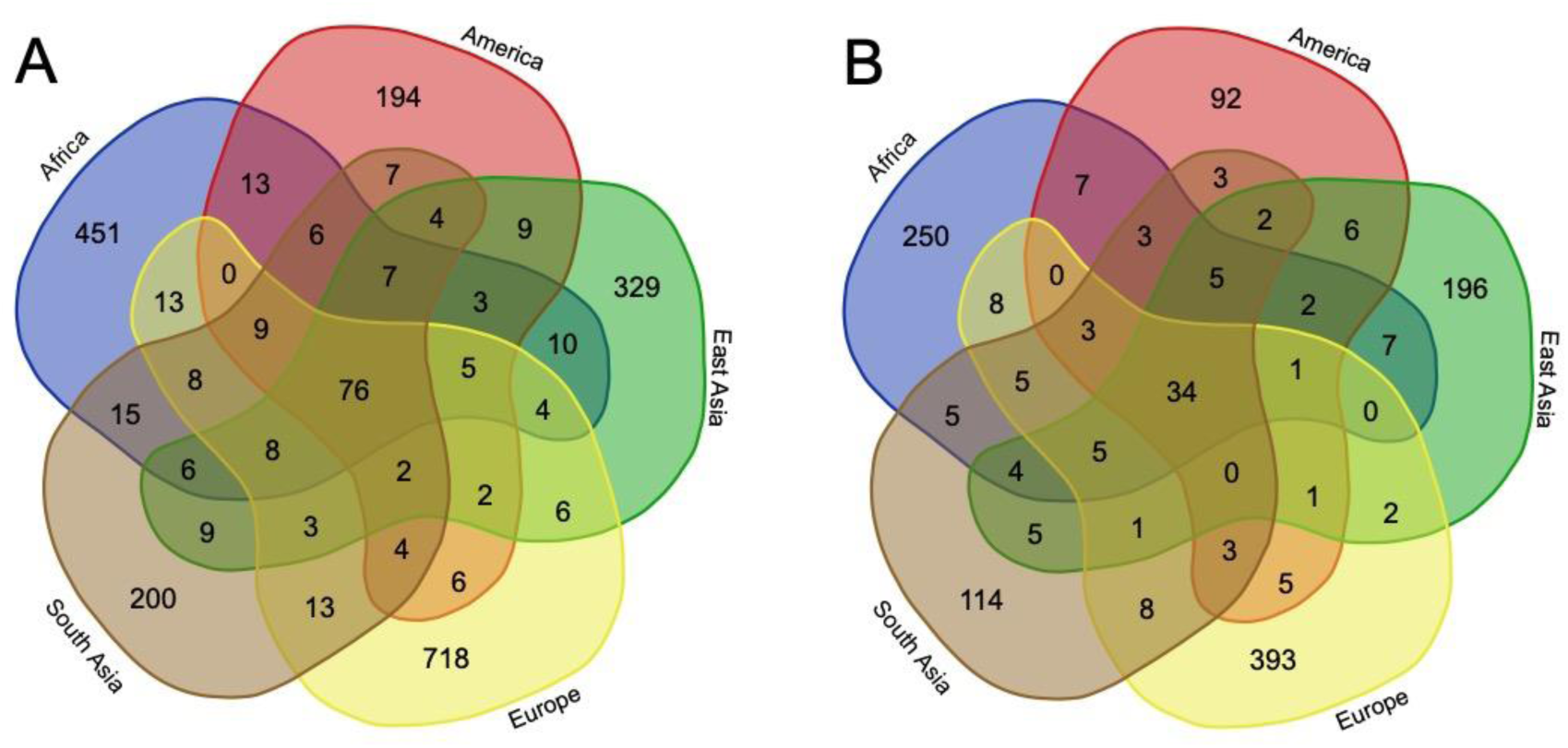
Venn diagrams of MIR sharing among super-populations. (A) Diagram of all MIRs shared among the five super-populations. (B) Diagram of only MIRs in gene regions according to the GENCODE database shared among the five super-populations.

In the current study, we defined an “MIR hit” as the number of MI alleles included within an MIR. Consequently, most MIRs in gene regions were unique in each super-population (i.e., singleton MIRs), especially the European population. There were a total of 1,008 singleton MIRs and 36 nonsingleton MIRs. The full list of MIR hit counts in gene regions among the super-populations is provided in Table S8. According to the results listed in Table S9, the most recurrent nonsingleton MIR in gene regions had eight hits in the African super-population. The second most recurrent MIR (hits = 7) was also detected in the African super-population.

### 2.5 Potential effects of MIs on human health

Although the MIs in normal individuals have not immediately impacted or directly caused diseases, MIs occurring in gene regions rather than intergenic regions are more likely to affect gene function and thus potentially result in phenotypic diversity in disease susceptibility among various populations in the future. To investigate the potential association of MIs with gene function and human diseases among various populations, we conducted an analysis of the genes that were frequently affected by MIs. We mainly focused on the eight genes that overlapped with MIRs (Table S9) with > 3 hits, as listed in Table S8. Then, we obtained gene functions through GeneCard as well as related reference studies; the functions are listed in Table S9. As shown in Table S9, MIs cause substitutions at the same position in gene regions of the hg19 human reference genome. Although it is not clear how these MIs in intronic or exonic (UTR) regions affect the functions of genes, it has been reported that both UTRs and exonic regions play an important role in translation and transcription [37,38]. In addition, most of the genes listed in Table S9 have been reported to be related to health. Among these genes, *ANKRD36* overlapped with MIs only in African populations. This gene is associated with genes that control the capacity to tolerate alcohol [39]. In addition to this MI, the MI located in MIR between chr10:61808458-61808486 caused an inverted sequence in one intron of Gene *ANK3*, potentially affecting an element region by which multiple sites are defined by ENCyclopedia of DNA Elements (ENCODE) [40] as being able to affect gene expression. According to the ENCODE definition, this MI is included in a CTCo transcription factor (To) binding site region between chr10:61807778-61808678, as derived from a large collection of ChIP-seq experiments performed by the ENCODE project [40], which could potentially affect the gene expression of *ANK3*. It has been reported previously that disease-related polymorphisms influence factor-binding site sequences in bipolar disease [41]. Additionally, SNPs correlated with disease susceptibility in bipolar disease have been recorded to alter the regulation of *ANK3* gene expression [42]. Considering the points above, the inverted allele between chr10:61808458-61808486, which causes 22 base substitutions in the intron region, may have some implications in terms of *ANK3* gene expression and function. In addition, *ANK3*, which overlapped with MIs in only East Asians, is reportedly a risk factor for schizophrenia, a chronic and severe mental disorder, in Han Chinese [43], which may explain the existence of MIs in *ANK3* only in East Asians.

### 2.6 Comparison with 7 nonhuman primates

Since we detected an important role of MIs in humans, it is worthwhile to explore the impact of MIs on the association of humans with nonhuman primates. In contrast to the individual genomes of the human populations from the 1KGP, herein, we used the alignments of human and 7 closely related nonhuman primate (chimpanzee, gorilla, orangutan, gibbon, baboon, rhesus macaque, and squirrel monkey) genomes to analyze MIs due to the lack of individual nonhuman primate genomes. Subsequently, we detected the MIs in the 7 nonhuman primates by applying the searchUMI tool [44] (see Materials and methods section for detection details) and identified a total of 24,476 MIs with the hg19 human assembly as a reference (Table S10). Notably, since MIRs were defined on the basis of the hg19 human genome assembly rather than the genome of the common ancestor of humans and the nonhuman primates, we adopted the MIs but not the MIRs for the nonhuman primates in this analysis.

Furthermore, we compared the MIs in humans with those in the 7 closely related nonhuman primates (chimpanzee, gorilla, orangutan, gibbon, baboon, rhesus macaque, and squirrel monkey) to investigate the inheritance and evolution of MI. By comparing inverted alleles in the genomes of nonhuman primates against the hg19 human reference genome and inverted alleles detected in the genomes of the 1KGP super-populations, we discovered a larger number of inverted alleles in African populations than in other populations that overlapped with those of the 7 nonhuman primates.

As shown in Table 2, the number of inverted alleles shared by nonhuman primates and the five super-populations in descending order were as follows: 58 shared with the African super-population, 49 with the East Asian super-population, 49 with the South Asian super-population, and 35 with the European super-population. The highest ratio of inverted allele overlap observed between the nonhuman primates and Africans might be because Africans inherited these shared inverted alleles from the common ancestor of nonhuman primates and humans which could be the ancestry alleles. Specifically, we speculate that after the Out of Africa geographic movement of modern humans, the African ancestral population spread the distribution of most ancestry alleles, and some of these ancestry alleles now survive in contemporary populations, resulting in the smaller inverted allele numbers in the other 4 super-populations. However, further sophisticated fossil and genetic data including the allele frequency based on more samples are necessary to support this conclusion.

**Table 2.** Counts of common inverted alleles among 7 nonhuman primates and five human super-populations.

| Name | All inverted allele count | East Asian | South Asian | European | African | American | 1KGP |
| --- | --- | --- | --- | --- | --- | --- | --- |
| Chimpanzee (panTro4) | 2,284 | 12 | 11 | 7 | 14 | 11 | 20 |
| Gorilla (gorGor3) | 2,010 | 10 | 11 | 9 | 14 | 10 | 18 |
| Orangutan (ponAbe2) | 3,473 | 9 | 9 | 7 | 10 | 9 | 14 |
| Gibbon (nomLeu1) | 3,713 | 6 | 7 | 5 | 7 | 3 | 13 |
| Baboon (PapHam1) | 4,547 | 5 | 5 | 3 | 6 | 5 | 8 |
| Rhesus macaque (rheMac3) | 4,271 | 5 | 4 | 2 | 5 | 4 | 7 |
| Squirrel monkey (SaiBol1) | 4,214 | 2 | 2 | 2 | 2 | 2 | 2 |
| Total | 24,476 | 49 | 49 | 35 | 58 | 44 | 82 |

## 3 Discussion and conclusions

In this study, we performed a comprehensive analysis of MIs in individual genomes from the 1KGP and 7 nonhuman primate genomes, built an MI landscape for the human populations, and explored the effect of MIs on human diversity, evolution, and health. Applying the software MID [13] for 26 human populations and searchUMI [44] for alignments between the human and 7 nonhuman primate genomes, we analyzed a total of 6,968 MIs detected in 1,937 individuals of 26 populations from the 1KGP and 82 inverted alleles common among the 7 nonhuman primate genomes and the 1,937 individuals. The MIs from the five super-populations (African, American, European, South Asian, and East Asian) revealed the population structure at multiple levels and may affect individuals in several ways. First, the widespread MI polymorphisms in human genomes greatly contribute to human diversity, which could be a result of geographical distribution and human migration. Second, large-scale MIs are good supplements to current evolutionary markers for reconstructing human evolutionary history. Third, the MIs in particular super-populations, especially the MIs overlapping with gene regions, are informative for understanding the association between genetic variations and health.

### 3.1 Average number of MIs in the five super-populations

We arranged the five super-populations in descending order based on the average number of inverted alleles per individual as follows: African > American > European > South Asian > East Asian (Fig. 3A). The average numbers of inverted alleles per individual among these five super-populations supported the Out of Africa hypothesis, which posits that humans originated in Africa and that recent Africans migrated to other continents. Other studies have speculated that modern Africans first migrated to Eurasia, which is the continent consisting of Europe and Asia [45]. In addition, the genetic distance between Africans and Europeans was reported to be smaller than that between Africans and Asians, which implies a closer relationship between Africans and Europeans [45]. In addition, as reported by Macaulay *et al* [46], the initial branch of the Asians from the southern dissemination followed the Nile from the east of Africa, went towards the north, and entered Asia. Once through the Sinai, the group of humans split, with some shifting into Europe and the others heading into Asia. This assumption is based on the comparatively late date of the landing of modern humans in Europe. Furthermore, the clustering result for the five super-populations based on MIs (Fig. 3B), which showed a tendency for European and Asian populations to cluster together and originate from Africans, also supported this hypothesis. In addition, East Asians were observed to have the greatest distance from Africans in the clustering results, which indicated that Asia was the last continent to which Africans arrived. It has been previously reported that during the migration to East Asia, a small number of Africans migrated along the Arabian Peninsula and Indian coast in South Asia and finally arrived in Australia [46]. The inverted allele number of Americans, which was between that of Africans and Europeans, is possibly due to both the African and European ancestry of the American populations. It is known that long-term migration can cause genomic variations [47]. Thus, we inferred that after the Out of Africa migration of modern human ancestors and hybridization among the four non-African super-populations, the Africans introduced some inverted alleles into the ancestors of the other four super-populations. It is speculated that the differential coevolution of MI lineages with different but closely related ancestral populations and subsequent evolution of MIs in parallel with the introgression of archaic alleles into the genomes of modern human ancestors may have been largely responsible for the present-day variant counts of MIs in multiple populations. Therefore, our results may provide evidence for the Out of Africa hypothesis.

### 3.2 Phylogenetic tree analysis and PCA

In the phylogenetic analysis of the 26 populations, geographical distributions and migrations affected the MIs. African and American populations were clustered on a branch in the phylogenetic tree, perhaps because African Americans are the largest racial minority [48] and mixed heritage is common in America. Specifically, the populations on the East Coast of America and in Western Africa were clustered on another branch. This cluster may be due to the black slave trade from Western Africa to the East Coast of America [49]. The populations on the West Coast of America and in Eastern Africa were more closely related than the other populations. This closeness is consistent with the migration of people from Eastern Africa to the West Coast of America [50]. We found that European and South Asian populations were clustered on one branch (as shown in Fig. 4), which has also been reported in other studies focusing on CNVs [51] and retroduplications [52]. The specificity of the Finnish population (FIN) can be interpreted in terms of multiple genetic components and demographic factors, such as isolation and migration, as reflected in their distinctive distribution among the European populations in another study [53]. Moreover, our finding that Puerto Ricans from Puerto Rico (PUR) are closely related to populations in East Asia is reasonable because some Peruvians are Asian immigrants [54]. These MIs used in phylogenetic tree analysis and PCA could also be used to cluster different super-populations, showing that these MIs represented the different genomic backgrounds of the studied populations. The results of the phylogenetic analysis of the 26 populations are consistent with the corresponding migration history and composition of these populations.

### 3.3 Comparative analysis with nonhuman primates

Evidence for the Out of Africa hypothesis also stems from a comparison between the MIs of nonhuman primates and those of humans. Africans shared more inverted alleles with the 7 nonhuman primates than did the other 4 super-populations, which shared only a few. The highest ratio of inverted alleles overlap observed between the nonhuman primates and Africans might be because Africans inherited these shared ancestry alleles from the common ancestor of nonhuman primates and humans in ancient times. It has been reported that genetic interchange among human populations, in terms of both periodic gene flow restricted by geographical space and events of main population expansion, contributed to interbreeding but not replacement [55]. Thus, it may be assumed that after the Out of Africa geographic movement of modern humans, the African ancestral population spread the distribution of ancestry alleles that are shared with the 7 nonhuman primates, and some of these ancestry alleles now survive in contemporary populations. Indeed, periodic genetic interchange occurred among human super-populations, in terms of both frequent gene flow constrained by geographical distance and population expansion events causing interbreeding. Because not all of the ancestry alleles in Africans would have been introduced into the other four super-populations, a smaller number of ancestry alleles remained in the other four super-populations, while some individuals finally inherited the derived allele. However, further sophisticated fossil and genetic data are needed to confirm this inference. MIs overlapped with 3 genes, *MYH14, GRM7*, and *DFNA5*, all of which are related to hearing [56-58], in both humans and nonhuman primates. This finding suggests that the inverted alleles in these 3 genes may have existed in the common ancestors of humans and nonhuman primates and were subsequently conserved in some nonhuman primates and some human super-populations. Therefore, the super-populations that inherited the inverted alleles may differ from others in terms of hearing.

Among the 76 common inverted alleles in the five super-populations, 34 overlapped with genes. The inverted alleles shared by all five super-populations could be a result of incompleteness of the current human reference assembly, while some of these regions could have been previously reported as SNPs because of the limited understanding of MIs. In addition, the human reference assembly has been reported to contain rare alleles, according to the Genome Reference Consortium. Accordingly, MIs are potential candidates in future updates of the human reference assembly.

### 3.4 Effects of MIs on human health

Here, we focused more on the effects of MIs on human evolution. However, as we analyzed the genes that overlapped with MIRs, we found that MIs may be associated with human health. Though the functions of the intronic regions and UTRs affected by the MIs detected with limited samples in this study are not yet clear, these MIs may benefit studies of the relation between variation and health in the future. In addition, the inverted alleles found in only 1 super-population, especially those overlapping with gene regions, may increase susceptibility to some diseases. Because of limited samples from normal individuals with no important genetic diseases in the 1KGP, the MIs in this study are candidates for future studies. Accordingly, more MIs from samples with diseases are needed to explore the correlation between MIs and human health. Nevertheless, we expect that personalized medical information will help us better understand the relationship between MIs and health, as well as unveil disease mechanisms, in the future.

## 4 Conclusions

To the best of our knowledge, this is the first study to analyze MIs in humans at the population scale. Our analysis of MIs with 1KGP data improves our understanding of human genetic diversity and evolution. The comparative analysis of MIs at the population, super-population, and species scales is thus expected to contribute to further implementation of human evolutionary theory. The future use of large-scale sophisticated fossil data and archeological materials to analyze the ages of MI polymorphisms will be informative in understanding the behavior of MIs as genetic markers and for better reconstructing human evolutionary history.

## 5 Materials and methods

### 5.1 Dataset

To elucidate the features of MIs among human genomes, we collected BAM files of the unmapped reads of 2,427 samples from the most recent release of the 1KGP [27]. Among all the samples, 490 did not contain MIs and were thus excluded. The 1,937 samples that included unmapped short reads with a total file size of 742 GB, which covered all 26 populations, were categorized into five super-populations: East Asians [Chinese Dai in Xishuangbanna, China (CDX); Han Chinese in Beijing, China (CHB); Southern Han Chinese (CHS); Japanese in Tokyo, Japan (JPT); and Kinh in Ho Chi Minh City, Vietnam (KHV)], South Asians [Bengali from Bangladesh (BEB); Gujarati Indians from Houston, Texas (GIH); Indian Telugu from the UK (ITU); Punjabi from Lahore, Pakistan (PJL); and Sri Lankan Tamil from the UK (STU)], Europeans [Utah Residents (CEPH) with Northern and Western Ancestry (CEU), Finnish in Finland (FIN), British in England and Scotland (GBR), Iberian Population in Spain (IBS), and Toscani in Italia (TSI)], Americans [Colombians from Medellin, Colombia (CLM); Mexican Ancestry from Los Angeles, USA (MXL); Peruvians from Lima, Peru (PEL); and Puerto Ricans from Puerto Rico (PUR)], and Africans [African Caribbeans in Barbados (ACB); Americans of African Ancestry in SW USA (ASW); Esan in Nigeria (ESN); Gambians in Western Divisions in the Gambia (GWD); Luhya in Webuye, Kenya (LWK); Mende in Sierra Leone (MSL); and Yoruba in Ibadan, Nigeria (YRI)]. The full list of all 1,937 sample names classified into 26 populations is shown in Table S1.

For further comparative analysis between humans and other primates, we also downloaded the hg19 human reference genome and the pairwise alignment data of 7 nonhuman primate assemblies from the UCSC Genome Browser (http://hgdownload.soe.ucsc.edu/downloads.html) [34]. We used data version panTro4 for the human/chimpanzee alignment, gorGor3 for the human/gorilla alignment, ponAbe2 for the human/orangutan alignment, nomLeu1 for the human/gibbon alignment, papHam1 for the human/baboon alignment, rheMac3 for the human/rhesus macaque alignment, and saiBol1 for the human/squirrel monkey alignment, all of which were downloaded from the UCSC Genome Browser (http://genome.ucsc.edu/).

### 5.2 MI detection and annotation

To detect MIs in data from the 1KGP, we applied the software MID [13] with default parameters and mapped all unmapped sequencing reads to the human reference genome (hg19) [34]. We chose MID because our previous investigation showed that MID was capable of efficiently identifying MIs from unmapped short reads through inversion reading and reference genome mapping. Specifically, the same MI occurring in multiple short reads of the same individual was counted as 1 MI during MID detection. After detecting MIs, we annotated them with both gene and correlated translated protein functions via the GENCODE database (https://www.gencodegenes.org/) [59], GeneCards database (http://www.genecards.org/) [60], and Metascape software [61]. The candidate promoter regions were defined as the regions within 2,000 bp upstream (for plus-strand genes) or downstream (for minus-strand genes) of gene regions. For the MIs overlapping with the CDS regions listed in Table S5, whenever possible we used the annotations for SVs and SNPs from the UCSC Genome Browser (http://genome.ucsc.edu/). The SVs and SNPs were annotated by the databases 1000G Ph3 Vars [27], dbSNP [31], ExAC [32], and ClinVar [33]. We detected the MIs of nonhuman primates in the aligned data using the searchUMI tool [44] because it is able to identify inversions ranging from 5 to 125 bp in length, which is the approximate length of MIs. During detection, the parameter pd was set at 0.0137 for panTro4, 0.0175 for gorGor3, 0.034 for ponAbe2, 0.029 for nomLeu1, 0.066 for papHam1, 0.065 for rheMac3, and 0.123 for saiBol1. Furthermore, based on the definition of MIs in this study, we removed inversions that were shorter than 10 bp.

### 5.3 Statistical analyses of MI diversity

Because sample number varied by population, diversity would be better reflected by the average number of inverted alleles per individual among super-populations or populations than by the total inverted allele number of each super-population or population. To address this point, we defined parameter 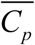 to represent the average count of inverted alleles per individual among super-populations or populations as follows: 

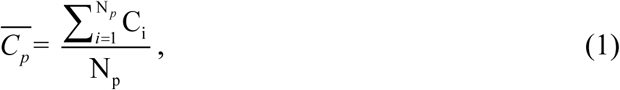

where C_i_ is the count of inverted alleles per individual, and Np is the total number of individuals in a population or super-population. The MI counts in each individual among the five super-populations, which were used to calculate the mean and SE of the inverted allele numbers per individual in the five super-populations, are listed in Table S11.

We used MIs but not MIRs to calculate the average inverted allele number because the MIR has merged a few MIs as one, and thus, MIR numbers cannot truly reflect the specific level of inverted allele numbers for each population. For the same reason, the phylogenetic tree and PCA analysis were built and performed based on the matrix of the MIs. However, only when analyzing the inverted alleles shared by the five super-populations with the Venn diagram did we adopt MIRs for analysis, because in this analysis, we expected to see overlapped alleles between the five super-populations and then to ascertain whether these overlapped regions had certain characteristics.

### 5.4 Population structure based on MIs

To analyze the population structure based on MIs among all populations from an evolutionary and comparative viewpoint, we next calculated a matrix with rows denoting MIRs and columns denoting all 26 human populations. Each element in this matrix represented the sum of inverted alleles included in the corresponding MIR within a given population. The matrices for constructing the phylogenetic tree based on all MIs are shown in Table S6. In addition, the matrices for constructing the phylogenetic tree based only on MIs in gene regions are provided in Table S7. With these matrices, we constructed a phylogenetic tree with the neighbor-joining algorithm via the R language. MIR sharing was analyzed by counting the number of shared MIRs among the five super-populations. The distances used in the neighbor-joining algorithm were Euclidean distances between every pair of column vectors, i.e., the distance between two populations was defined as the average MIR distance between pairs of individuals from the two populations. In the same way, the phylogenetic tree of the five super-populations was also constructed based on the neighbor-joining clustering method using the MIs obtained in gene regions (Table S7).

MIR sharing was analyzed by counting the number of shared MIRs among the five super-populations. The population structure of the 26 populations was also visualized through PCA with the R programming language to determine whether the distribution of MIs was based on geographical location and migration history. For the PCA, we applied the same matrix (Table S6) used in the phylogenetic tree analysis. In addition, we regarded the 2,140 MIRs as the principal components in the PCA.

