## Supplementary_Material for "The landscape of micro-inversions provides clues for population genetic analysis of humans"

Supplemental Material includes two figures and ten tables. Supplementary Tables are included as worksheets in the files Table S1.xlsx ~ Table10.xlsx

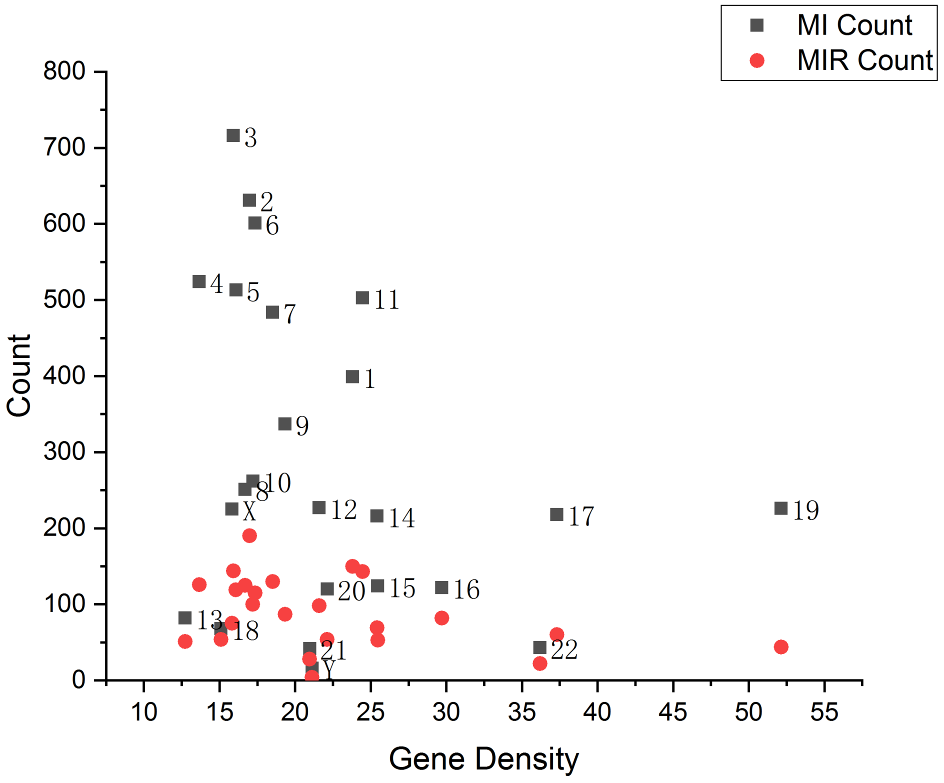


**Figure S1. Scatter plot of counts of MI and MIR by gene density.** We discovered that chromosomes 19, 17, 16, and 11 had high gene density.

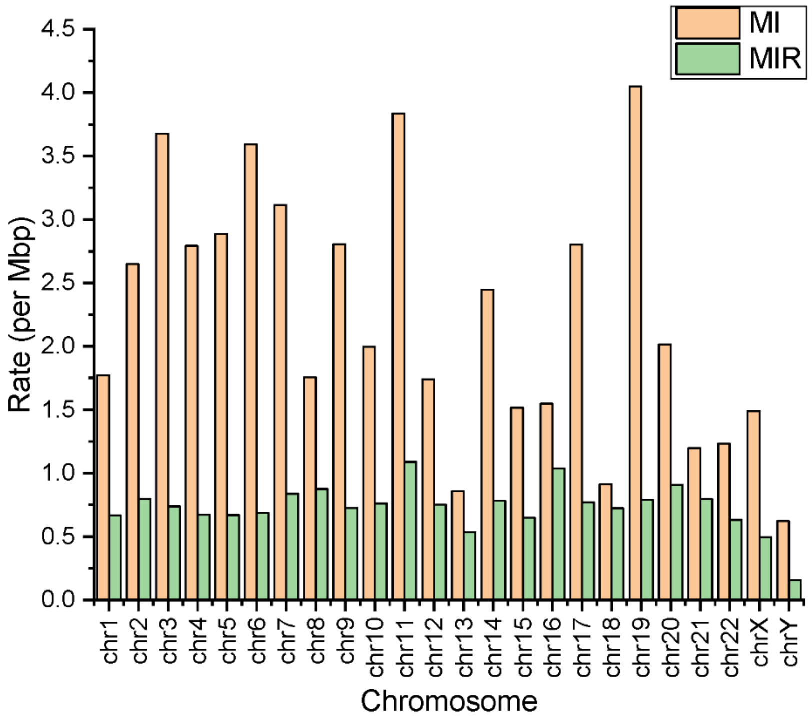


**Figure S2. Distribution of the MI and MIR event rate across chromosomes.**

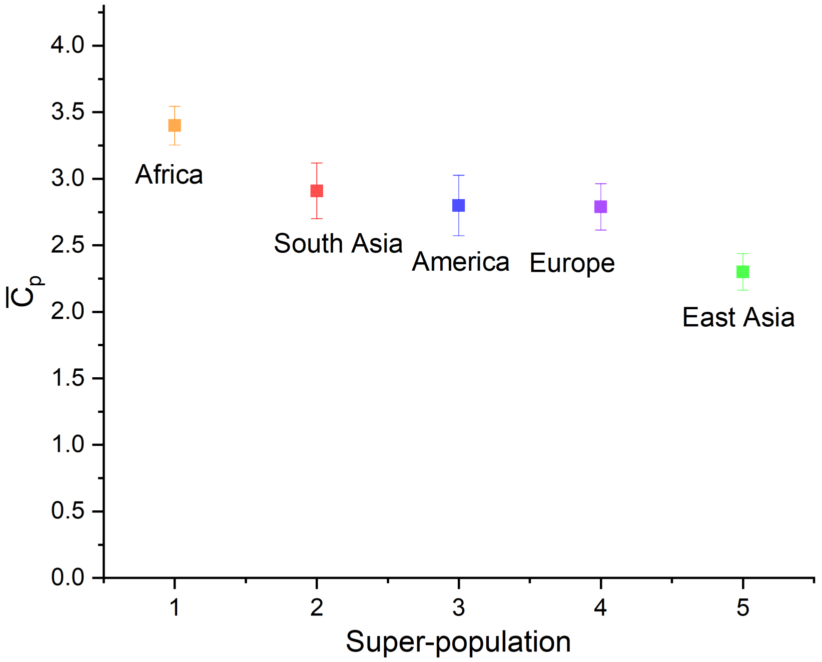


**Figure S3. Average counts of inverted alleles per individual for five super-populations taking account into samples with no inverted alleles.**

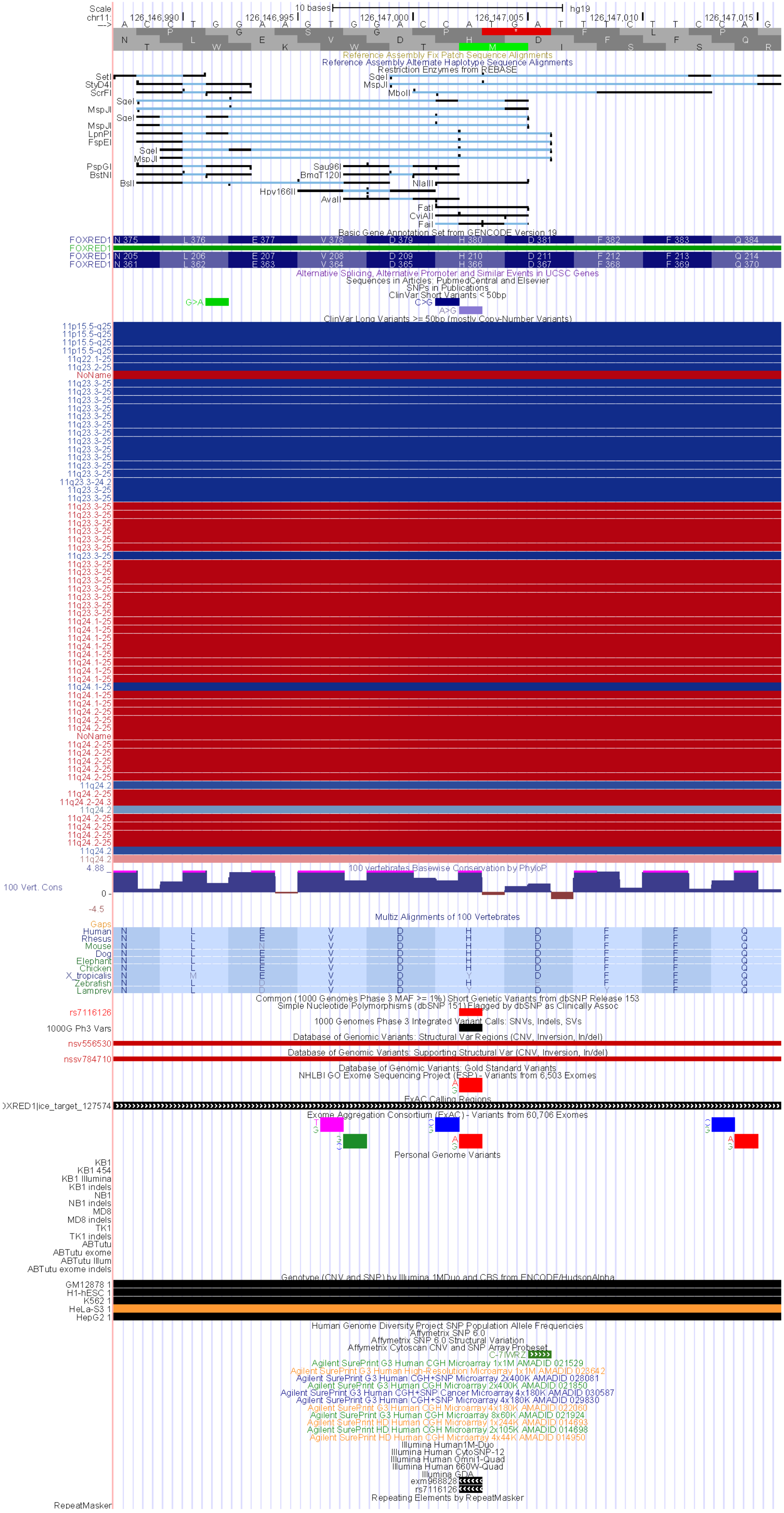


**Figure S4. Annotation of MI chr11: 126146988- 126147016 from the UCSC Genome Browser[1].**

### Supplemental Tables (in separate spreadsheets)

**Table S1. The list of 1,937 samples from the 1KGP, grouped by populations.** Table S1 lists all 1,937 samples downloaded from the 1KGP in our study. The 1,937 samples were grouped by 26 populations. In summary, there are 81 samples in ACB, 44 samples in ASW, 68 samples in BEB, 62 samples in CDX, 80 samples in CDU, 80 samples in CHB, 80 samples in CHS, 70 samples in CLM, 87 samples in ESN, 77 samples in FIN, 74 samples in GBR, 83 samples in GIH, 102 samples in GWD, 70 samples in IBS, 86 samples in ITU, 90 samples in JPT, 74 samples in KHV, 83 samples in LWK, 81 samples in MSL, 55 samples in MXL, 63 samples in PEL, 85 samples in PJL, 35 samples in PUR, 91 samples in STU, 75 samples in TSI, and 50 samples in YRI.

**Table S2. MIs and annotations detected from the 1KGP data.** Table S2 lists all 6,968 MIs detected in our study. From left to right, the column shows the sequence read name in which we detected MIs, the chromosome in which the MI is located, start of coordinate MIs, end of coordinate MIs, MI length, MI sequence, whether MI is in the gene or intergenic region, gene name overlapping with MI, gene type overlapping with MI, status of gene overlapping with MI, exon count overlapping with MI, CDS region count overlapping with MI, and UTR count overlapping with MI.

**Table S3. MIRs and annotation.** Table S3 lists all 2,140 MIRs in our study. From left to right, the column shows the MIR serial numbers, chromosome in which MIR is located, start of coordinate MIRs, end of coordinate MIRs, MIR length, counts for MIs located in the MIR, whether MIR is in the gene or intergenic region, gene name overlapping with MIR, gene type overlapping with MIR, status of gene overlapping with MIR, exon count overlapping with MIR, CDS region count overlapping with MIR, UTR count overlapping with MI.

**Table S4. MIs overlapping with exon regions.** Table S4 lists all MIs overlapping with exon regions.

**Table S5. MIs overlapping with CDS regions.** Table S5 lists all the MIs overlapping with CDS regions. The annotations for SVs and SNPs are from the UCSC Genome Browser (http://genome.ucsc.edu/). The SVs and SNPs were annotated by the database of 1000G Ph3 Vars including 1000 Genomes Phase 3 Integrated Variant Calls: SNVs, Indels, SVs, dbSNP, ExAC, ClinVar, and Database of Genomic Variants: Structural Var Regions.

**Table S6. Matrix for all MIs in the phylogenetic tree and PCA analysis.** Table S5 lists the matrix data of all 6,968 MIs used for the phylogenetic tree and PCA analysis, with rows denoting the 2,140 MIRs and columns denoting all 26 human populations. Each element in this matrix represents the sum of MIs included in the corresponding MIR within this population.

**Table S7. Matrix for MIs in gene regions for phylogenetic tree analysis.** Table S6 lists the matrix data for the 2,135 MIs in gene regions used for phylogenetic and PCA analyses, with rows denoting the 2,140 MIRs and columns denoting all 26 human populations and seven non-human primates. Each element in this matrix represents the sum of MIs included in the corresponding MIR within this population.

**Table S8. MIR hits in gene regions in five super-populations.** Table S7 lists the counts for MIs supported by multiple samples in gene regions in five super-populations. Numbers in brackets represent the MIR hit counts in gene regions in five super-populations. For example, 321 MIs in the gene region in East Asia were supported by only one and six MIs in the gene region in East Asia were supported by two samples.

**Table S9. Population-specific genes overlapping with MIRs.** Table S8 lists ten genes overlapping with MIs that were only present in one super-population. The influence on health and function of the ten genes is also listed in Table S8.

**Table S10. Shared MIs among different non-human primates.** Table S9 lists shared MIs among six non-human primates. From left to right, each column represents names of primates sharing MIs, the number of MIs shared by non-human primates, sequence read names in which MIs are located, whether MIs are in gene or intergenic regions, gene names where MI are located, whether the gene is protein-coding, and whether the gene is known.

**Table S11. MI number in each individual among five super-populations.** Table S10 lists the MI number for each individual used to calculate the means and standard errors of MI numbers among five super-populations. The rows "Mean" and "Standard Error" represent the average number and standard error of the MI number in each super-population, respectively.

### Supplemental Figures:

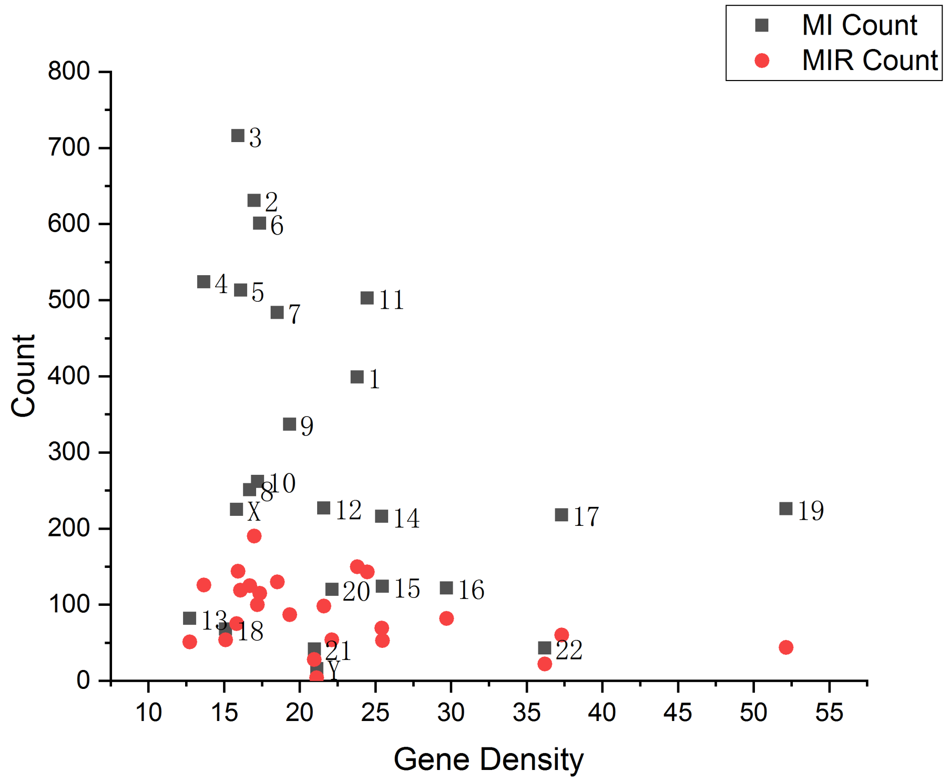


**Figure S1. Scatter plot of counts of MI and MIR by gene density.** We discovered that chromosomes 19, 17, 16, and 11 had high gene density.

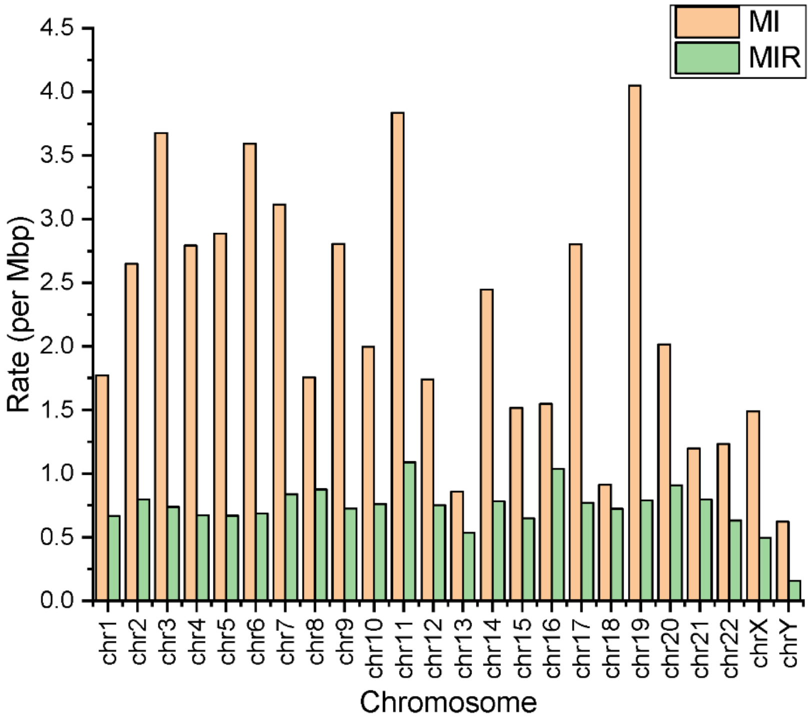


**Figure S2. Distribution of the MI and MIR event rate across chromosomes.**

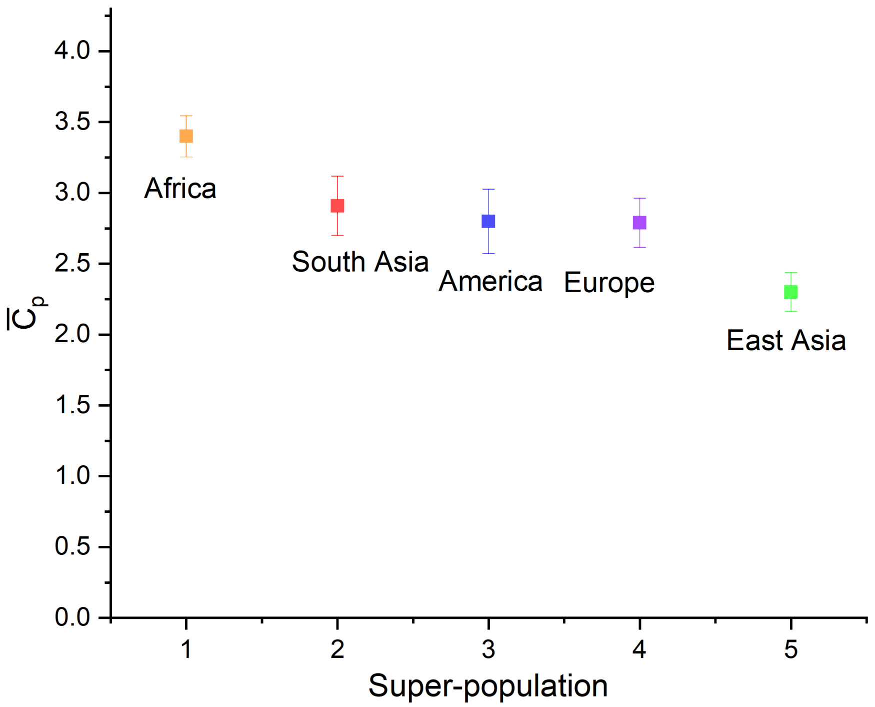


**Figure S3. Average counts of inverted alleles per individual for five super-populations taking account into samples with no inverted alleles.**

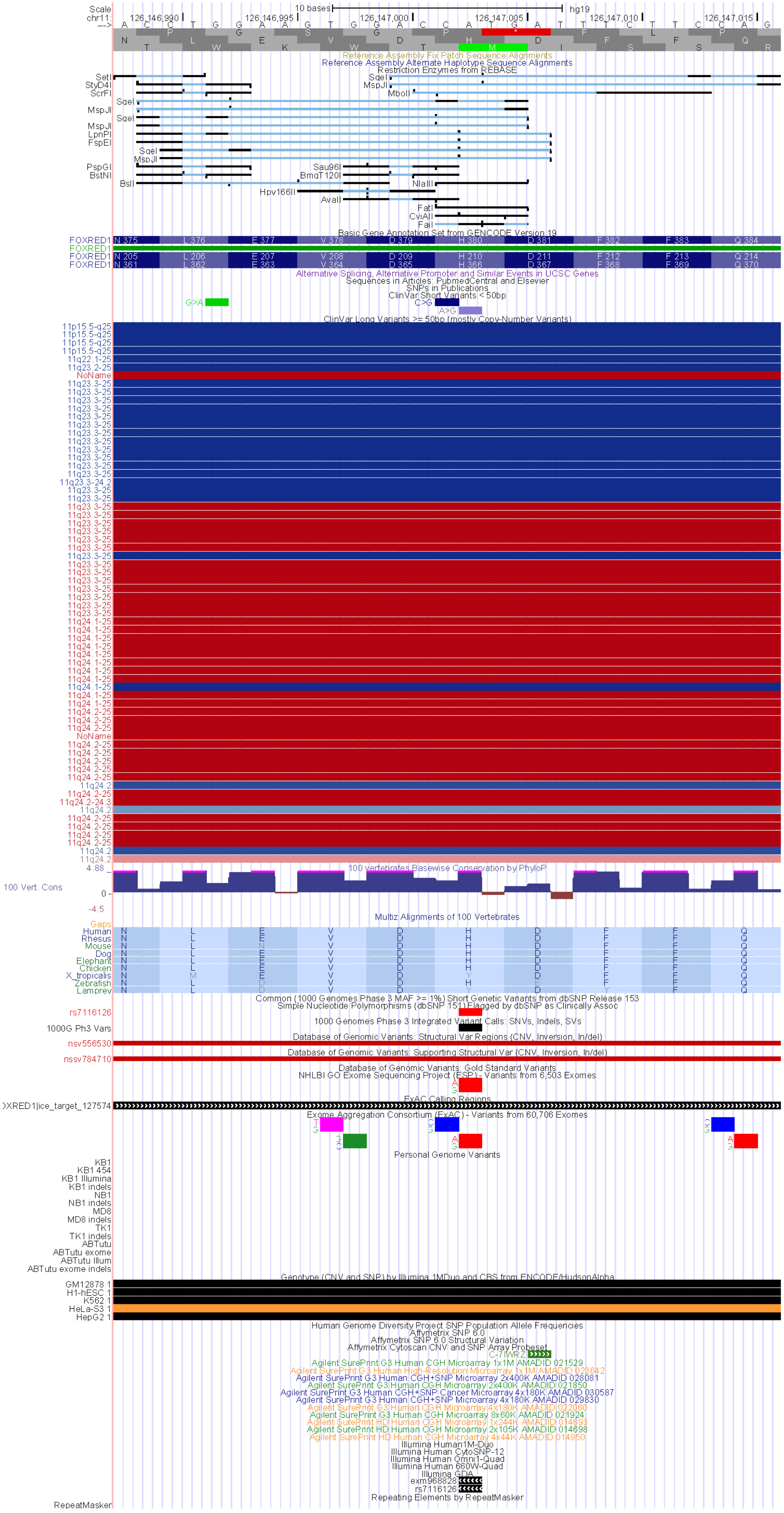

**Figure S4. Annotation of MI chr11: 126146988- 126147016 from the UCSC Genome Browser[1].**
